# The oncogenic Kaposi’s sarcoma-associated herpesvirus encodes a mimic of the tumor suppressive miR-15/16 miRNA family

**DOI:** 10.1101/660811

**Authors:** Kylee Morrison, Mark Manzano, Kevin Chung, Matthew J. Schipma, Elizabeth T. Bartom, Eva Gottwein

## Abstract

Many tumor viruses encode oncogenes of cellular origin. Here, we report the first instance of an oncoviral mimic of a cellular tumor suppressor. The Kaposi’s sarcoma-associated herpesvirus (KSHV) miRNA miR-K6-5p shares sequence similarity to the tumor suppressive cellular miR-15/16 miRNA family. We show that miR-K6-5p inhibits cell cycle progression, a hallmark function of miR-16. miR-K6-5p regulates conserved miR-15/16 targets, including many cell cycle regulators. Inhibition of miR-K6-5p in KSHV-transformed B cells conferred a significant growth advantage. Altogether, our data show that KSHV encodes a functional mimic of the miR-15/16 miRNA family. While it is exceedingly well established that oncogenic viruses encode oncogenes of cellular origin, this is the first example of an oncogenic virus that encodes a viral mimic of a cellular tumor suppressor. Encoding a tumor suppressive miRNA could help KSHV balance viral oncogene expression and thereby avoid severe pathogenesis in the healthy host.

## Introduction

Viruses cause ∼12% of human cancers. Viral oncogenesis is often due to the expression of viral oncogenes, including those with cellular counterparts. Kaposi’s sarcoma-associated herpesvirus (KSHV) is a human tumor virus that causes Kaposi’s sarcoma (KS) and primary effusion lymphoma (PEL), and the B cell proliferative disorder multicentric Castleman’s disease (Cesarman et al., 1995; Chang et al., 1994; Nador et al., 1996; Soulier et al., 1995). The vast majority of cells in KS and PEL exhibit the restricted latent KSHV gene expression pattern, which includes three proteins that promote proliferation and survival of infected cells: LANA maintains the viral episome and inhibits the tumor suppressor p53 (Ballestas et al., 1999; Friborg et al., 1999), the KSHV cyclin (vCyc) drives cell cycle progression (Chang et al., 1996; Li et al., 1997), and the KSHV FLICE-inhibitory protein (vFLIP) promotes cellular survival (Guasparri et al., 2004). The latency program also includes >20 viral microRNAs (miRNAs) that have the potential to substantially reshape host gene expression (Cai et al., 2005; Grundhoff et al., 2006; Pfeffer et al., 2005; Samols et al., 2005).

miRNAs are ∼22 nucleotide (nt) long non-coding RNAs that associate with Argonaute proteins in RNA-induced silencing complexes (RISCs) to mediate mRNA repression (Bartel, 2009, 2018). The large majority of effective miRNA binding sites exhibit uninterrupted Watson-Crick base pairing to nts 2-7 from the miRNA 5’end, the seed sequence. An effective regulatory outcome is achieved by additional base pairing of nt 8 of the miRNA or the presence of an adenosine (A) immediately following the seed match in the target mRNA. miRNA-mediated target regulation results in measurable target mRNA destabilization (Bagga et al., 2005; Jonas and Izaurralde, 2015; Lim et al., 2005).

Together, the KSHV miRNAs bind hundreds of mRNAs and therefore have a pleiotropic functional outcome (Gallaher et al., 2013; Gay et al., 2018; Gottwein et al., 2011; Grosswendt et al., 2014; Haecker et al., 2012; Ziegelbauer et al., 2009). Roles of individual KSHV miRNAs include the evasion from cell cycle arrest and apoptosis (Gottwein and Cullen, 2010; Liu et al., 2017). Interestingly, KSHV uses at least three viral miRNAs to access conserved cellular miRNA regulatory networks (Gottwein et al., 2011; Gottwein et al., 2007; Manzano et al., 2015; Manzano et al., 2013; Skalsky et al., 2007). Most importantly, miR-K11 shares its seed sequence with the oncogenic miRNA miR-155 and consequently regulates miR-155 targets. miR-K11 phenocopies miR-155-induced B cell proliferation *in vivo* (Boss et al., 2011; Dahlke et al., 2012; Sin et al., 2013). miR-K11 is therefore likely to contribute to KSHV-associated B cell lymphomagenesis.

Interestingly, the KSHV miRNA miR-K6-5p has extended sequence similarity to the cellular miR-15/16 family of miRNAs (Fig. 1A). This is surprising, because miR-15/16 family miRNAs are tumor suppressors. miR-15/16 family miRNAs are encoded within several miRNA clusters. Chromosomal deletion of 13q14, which harbors the miR-15a/miR-16-1 cluster, is frequent in B-cell chronic lymphocytic leukemia (CLL) and can result in substantially reduced miR-15/16 expression (Calin et al., 2002; Fulci et al., 2007; Wang et al., 2008). Downregulation of miR-15/16 has also been reported in other cancers. In mouse models, individual or combined deletion of the miR-15a/miR-16-1 and miR-15b/miR-16-2 clusters leads to hematologic malignancies (Klein et al., 2010; Lovat et al., 2015; Lovat et al., 2018). At the cellular level, miR-15/16 inhibit cell cycle progression and promote apoptosis and targets of miR-15/16 include strong candidates for mediators of its tumor suppressive properties, such as pro-apoptotic proteins of the BCL2 family (Cimmino et al., 2005), several cyclins, cyclin-dependent kinases, and other cell cycle regulators (Calin et al., 2008; Linsley et al., 2007; Liu et al., 2008).

**Figure 1.**
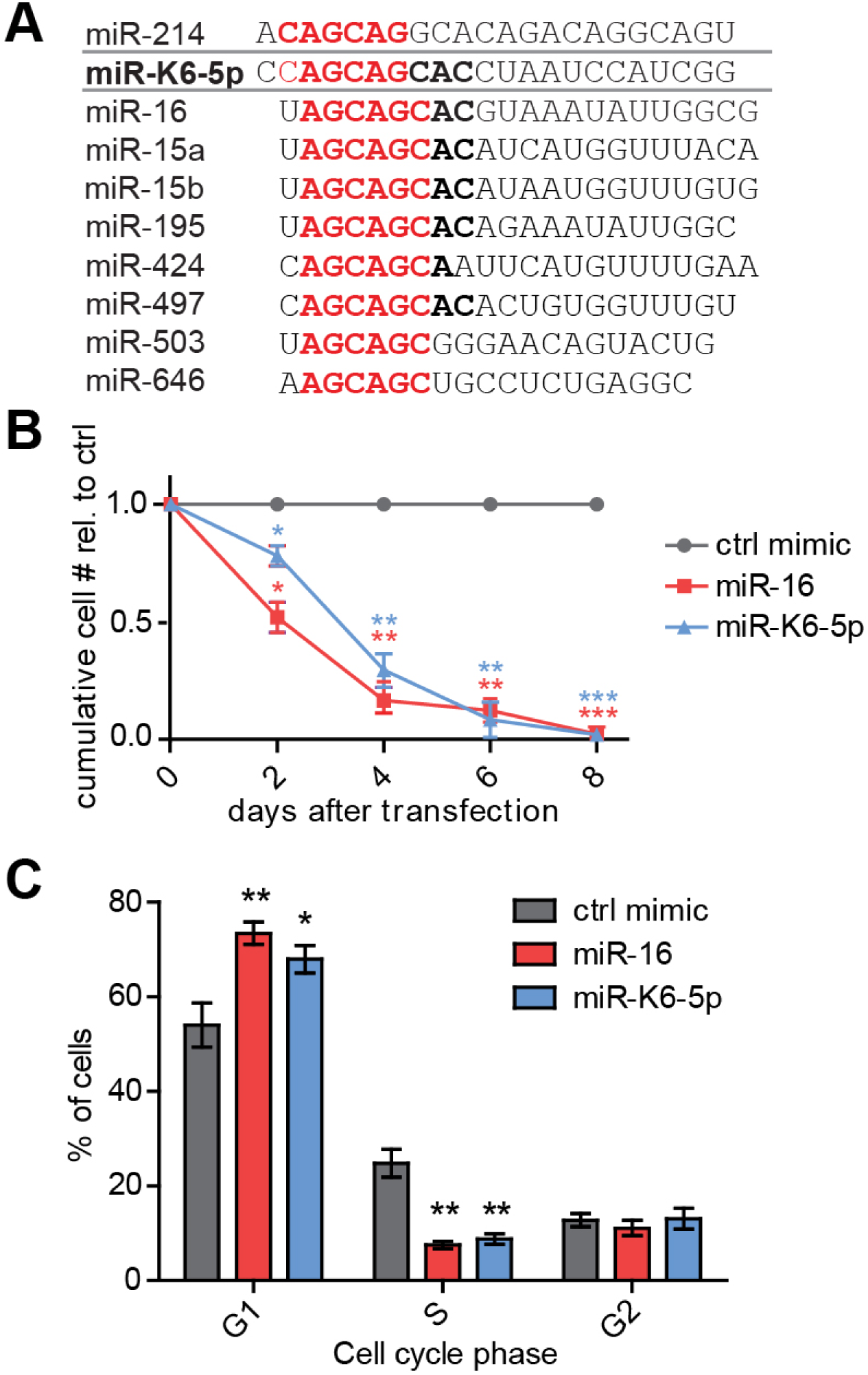
KSHV miR-K6-5p shares sequence similarity with the miR-15/16 family and mimics miR-16-induced cell cycle arrest. (A) Sequences of miR-K6-5p, the cellular miR-15/16 miRNA family, and miR-214. nts 2-7, comprising the minimal seed sequence of each miRNA, are in red. nts that contribute to the extended similarity between miR-K6-5p and miR-15/16 are in bold. (B) Primary lymphatic endothelial cells (LEC) were transfected with 10nM miRNA mimics of miR-16, miR-K6-5p, or a negative control (ctrl) mimic, counted and split to equal numbers every 2 days. Cell numbers were normalized to those of ctrl-transfected cells. (n=3, * p<0.05, ** p<0.01, *** p<0.001, error bars represent s.e.m). (C) LEC were transfected as described for (B). 2 days after transfection, cells were stained with propidium iodide (PI) to measure DNA content and analyzed by FACS. (n=3, * p<0.05, ** p<0.01, error bars represent s.e.m). Also see Fig. S1.

Here, we characterize the functional relationship between KSHV miR-K6-5p and the miR-15/16 family miRNAs. miR-K6-5p phenocopies the cell cycle inhibitory functions of miR-15/16, shares a substantial portion of its mRNA target sites, and inactivation of miR-K6-5p confers a competitive advantage to KSHV-transformed B cells. Thus, this oncogenic virus remarkably encodes a mimic of a conserved tumor suppressor. We hypothesize that the physiological role of miR-K6-5p is to negatively regulate the cell cycle and thereby balance the pro-proliferative and pro-survival functions of the KSHV oncogenes. While the expression of tumor-viral versions of cellular oncogenes is a well-established paradigm, miR-K6-5p is the first documented oncoviral mimic of a *bona fide* cellular tumor suppressor.

## Results

### miR-K6-5p mimics the cell cycle inhibitory properties of miR-16

To investigate the relationship between miR-K6-5p and miR-16, we compared consequences of miRNA mimic transfection into primary lymphatic endothelial cells (LECs). LECs are candidates for the cell type of origin of KS and a commonly used model system for the study of KSHV (Boshoff et al., 1995; Staskus et al., 1997; Wang et al., 2004). miR-16 was chosen as a representative of the miR-15/16 family members based on its high expression in PEL cell lines (Gottwein et al., 2011). Other family members were not included, because miR-15, miR-16 and miR-195 have previously been shown to induce nearly identical changes in gene expression and phenotypes (Linsley et al., 2007). As expected, overexpression of miR-16 strongly reduced live cell numbers over time (Fig. 1B) and arrested cells at the G1 phase of the cell cycle (Fig. 1C, Fig. S1A). Interestingly, expression of miR-K6-5p similarly reduced proliferation and caused cell cycle arrest at G1. miR-15/16 are ubiquitously expressed, which makes their direct comparison to miR-K6-5p difficult. To enable a more direct comparison, we deleted both miR-15/16 clusters from 293T cells (293T/DKO). A resulting 293T/DKO cell clone was transduced with an empty lentiviral vector, a miR-K6-5p expression vector (Fig. S1B), or a miR-16 expression vector. Primer extension analysis confirmed a complete absence of endogenous miR-16 expression in 293T/DKO and demonstrated re-expression of physiological levels of miR-16 upon transduction with the miR-16 expression vector (Fig. S1C). Primer extension analysis furthermore confirmed that ectopically expressed miR-K6-5p exhibits an accurate 5’-end and expression levels comparable to those observed in the PEL cell line BC-3. Growth curve analysis showed that miR-K6-5p and miR-16 similarly reduce live cell 293T/DKO numbers over time (Fig. S1D). Together, these results demonstrate that miR-K6-5p has cell cycle inhibitory properties similar to miR-16.

### miR-K6-5p mimics miR-16-induced effects on mRNA expression

We next used mRNA sequencing (mRNA-Seq) to determine whether the observed phenotypic mimicry between miR-K6-5p and miR-16 is due to functional mimicry at the level of mRNA regulation. This experiment was performed in 293T cells lacking Dicer (293T/NoDice) (Bogerd et al., 2014), which do not express any miR-15/16 family miRNAs and therefore provide a clean background for a direct comparison between viral miR-K6-5p and the cellular miRNAs it resembles. Both miR-16 and miR-K6-5p caused cell cycle arrest in 293T/NoDice cells compared to a negative control mimic (Fig. S1E-F). Interestingly, miR-16 transfected 293T/NoDice cells showed a G1 arrest, while miR-K6-5p-transfected cells may instead be arrested at G2, hinting at functional differences between these miRNAs that affect the cell cycle in a cell-type dependent manner. For mRNA-Seq, 293T/NoDice cells were transfected with mimics of miR-16, miR-K6-5p, a variant of miR-K6-5p with a 5’-terminal C to U substitution (miR-K6-5p-5’U), miR-214, or a control mimic. miR-K6-5p-5’U was included to control for effects that result from the 5’-C of miR-K6-5p, which is rarely seen in cellular miRNAs and may lead to suboptimal loading into RISC (Seitz et al., 2011). Since the miRNA 5’-nucleotide does not participate in target selection, this substitution is not expected to alter the range of regulated mRNAs. Finally, miR-214 was included, because this miRNA shares its exact seed region (nts 2-7) with miR-K6-5p (Fig. 1A). miR-214 is a poorly conserved cellular miRNA of largely uncharacterized function.

Principal component analysis of the resulting dataset revealed that, despite the offset seed region, gene expression changes in miR-K6-5p-transfected cells were more similar to miR-16-transfected cells than to miR-214-transfected cells (Fig. 2A, Table S1). Results from miR-K6-5p-5’U were indistinguishable to miR-K6-5p, showing that potential effects of the 5’C in WT miR-K6-5p do not confound the results under our experimental settings (Figs. 2A, C; S2A). Direct pairwise comparison of mRNA expression changes induced by miR-16 and miR-K6-5p detected a highly significant correlation, strongly suggesting that miR-16 and miR-K6-5p affect gene expression similarly (Fig. 2B-C). In contrast, correlation of either miR-16 or miR-K6-5p-induced changes in mRNA expression with those resulting from miR-214 expression was considerably more modest (Figs. 2C; S2B-C). Gene Set Enrichment Analysis (GSEA) showed that miR-16-and miR-K6-5p-downregulated genes were similarly enriched in functional categories broadly associated with cellular proliferation (Fig. S2D) (Subramanian et al., 2005). Together, these results suggest that miR-K6-5p-induced mRNA expression changes are more similar to those induced by miR-16, whose seed is offset by one nt, than by miR-214, which shares its exact hexamer seed sequence (nts 2-7) with miR-K6-5p.

**Figure 2.**
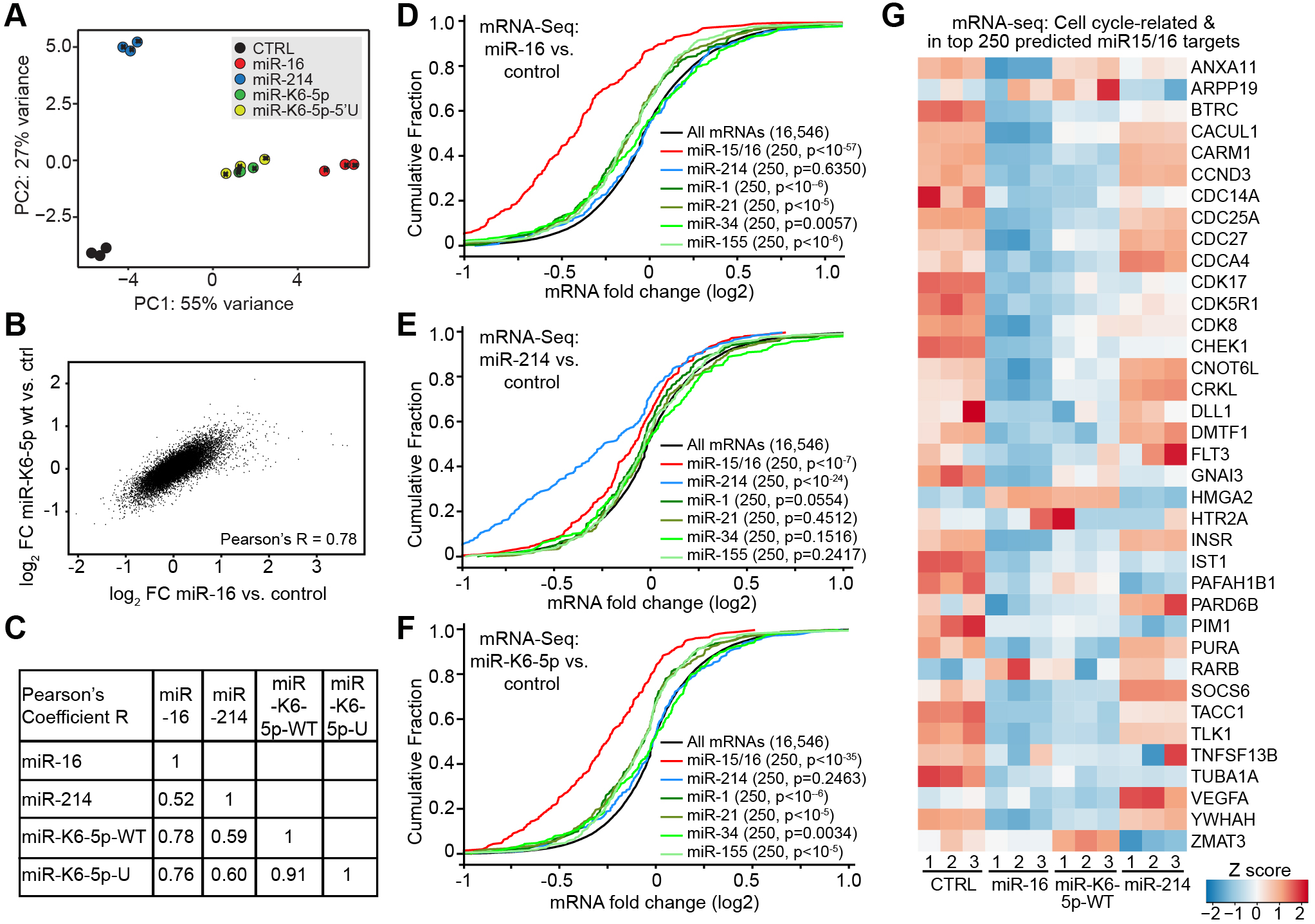
miR-K6-5p mimics miR-16-induced gene expression changes. (A) 293T/NoDice were subjected to mRNA-Seq two days after transfection with miRNA mimics. Principle component (PC) analysis of mRNA-Seq data. (B) Pearson Correlation was used to compare mRNA log2 fold changes in miR-16 and miR-K6-5p transfected cells. (C) Table of Pearson’s Coefficients from additional comparisons in the mRNA-Seq dataset, as in B (see Figure S2A-C). (D-F) Cumulative distribution frequency (CDF) plots depicting regulation of the top 250 TargetScan-predicted targets of the listed miRNAs by mimics of miR-16 (D), miR-214 (E), or miR-K6-5p-WT (F) in the mRNA-Seq experiment. Numbers in parentheses indicate gene set sizes and *p* values for comparisons to all mRNAs, which were calculated using 2-sample K-S tests. (G) Heatmap showing Z scores for mRNAs among the top 250 TargetScan-predicted miR-15/16 targets that contribute to the enrichment in cell cycle-related categories as detected by DAVID (see Table S2). Also see Supplementary Fig. 2 for additional analyses of the mRNA-Seq dataset and Table S1 for gene sets.

### miR-K6-5p downregulates miR-16 targets

We next tested whether miR-K6-5p regulates miR-16 targets. To establish a set of high confidence targets of miR-16, we used our mRNA-Seq dataset to compare miR-16-mediated regulation of mRNAs that were computationally predicted as miR-15/16 family targets by Targetscan or experimentally identified as candidates for miR-16 targets by the crosslinking-based methods PAR-CLIP or CLASH (Agarwal et al., 2015; Gay et al., 2018; Gottwein et al., 2011). While candidates for miR-16 targets from each approach were significantly repressed by miR-16, the top 250 Targetscan-predicted targets were most strongly regulated and therefore used for further analysis (Fig. S2E). Targetscan predicts and ranks mRNA targets of cellular miRNAs based on the number and types of seed matches, evolutionary conservation and sequence context (Agarwal et al., 2015). The observed regulation of the predicted miR-15/16 family targets by miR-16 was specific, because similarly predicted targets of miR-214, the tumor suppressive miRNA miR-34a, the tissue-specific miR-1, or the oncomiRs miR-155 or miR-21 were not substantially affected by miR-16 (Fig. 2D). miR-214 significantly regulated the top 250 predicted targets of miR-214, but only weakly downregulated predicted targets of miR-15/16 (Fig. 2E). Strikingly, miR-K6-5p strongly and specifically downregulated miR-16 targets (Fig. 2F). In contrast, miR-K6-5p failed to repress predicted targets of miR-214, despite sharing its hexamer seed with miR-214 (Fig. 2F). The top 250 predicted miR-16 targets were highly enriched for cell cycle regulators (Table S2). The large majority of the mRNAs encoding these cell cycle regulators was repressed by both miR-16 and miR-K6-5p, but not miR-214 (Fig. 2G). Therefore, functional mimicry of miR-16 by miR-K6-5p extends to the repression of mRNAs encoding cell cycle regulators, including, for example, the known miR-16 targets CCND3 (Liu et al., 2008) and CDC25A (Pothof et al., 2009). We obtained overall similar results by analyses of mRNAs based simply on the presence of canonical miRNA binding sites within their 3’UTRs (Fig. S2F-I). Together, these results further support the conclusion that miR-K6-5p is functionally closer to miR-16 than to miR-214. These results also demonstrate that miR-K6-5p specifically represses a large subset of the evolutionarily conserved targets of miR-15/16, including many cell cycle regulators.

### miR-K6-5p shares binding sites with miR-16

We next tested whether miR-K6-5p regulates miR-16 targets through shared binding sites. CLASH (crosslinking, ligation, and sequencing of hybrids, Helwak et al., 2013) employs ligation reactions that allow the direct identification of miRNA binding sites, without the need for computational assignment of the targeting miRNA. A published CLASH dataset from an endothelial model of KSHV infection (Gay et al., 2018) identified 18 3’UTR hybrids with perfect seed base pairing to miR-K6-5p. Of these sites, 11 (61%) were also ligated to miR-16, which offers direct evidence that miR-K6-5p shares a substantial portion of its binding sites with miR-16 (Table S3). A closer look at the sites identified by qCLASH and the miR-16 targets that are repressed by miR-K6-5p in our mRNA-Seq experiment reveals that the unusual trinucleotide repeat seed sequences of miR-16 and miR-K6-5p enable a substantial overlap in binding sites (Tables S1, 3). Specifically, seed matches to nts 2-8 of miR-16 that are immediately flanked by a 3’-G (“miR-16 2-8G”) or preceded by 5’-GC (“miR-16 GC2-8”) are canonical miRNA binding sites for miR-K6-5p (Fig. 3A). Together, these sites are surprisingly abundant. For example, ∼51% of all mRNAs with 2-8mer seed matches to miR-16 in the mRNA-Seq dataset contain 2-8G and/or GC2-8 sites for miR-K6-5p. However, our analyses also suggest that both miR-K6-5p and miR-16 have some targets that are not expected to be shared (Table S1, Fig. 3A).

**Figure 3.**
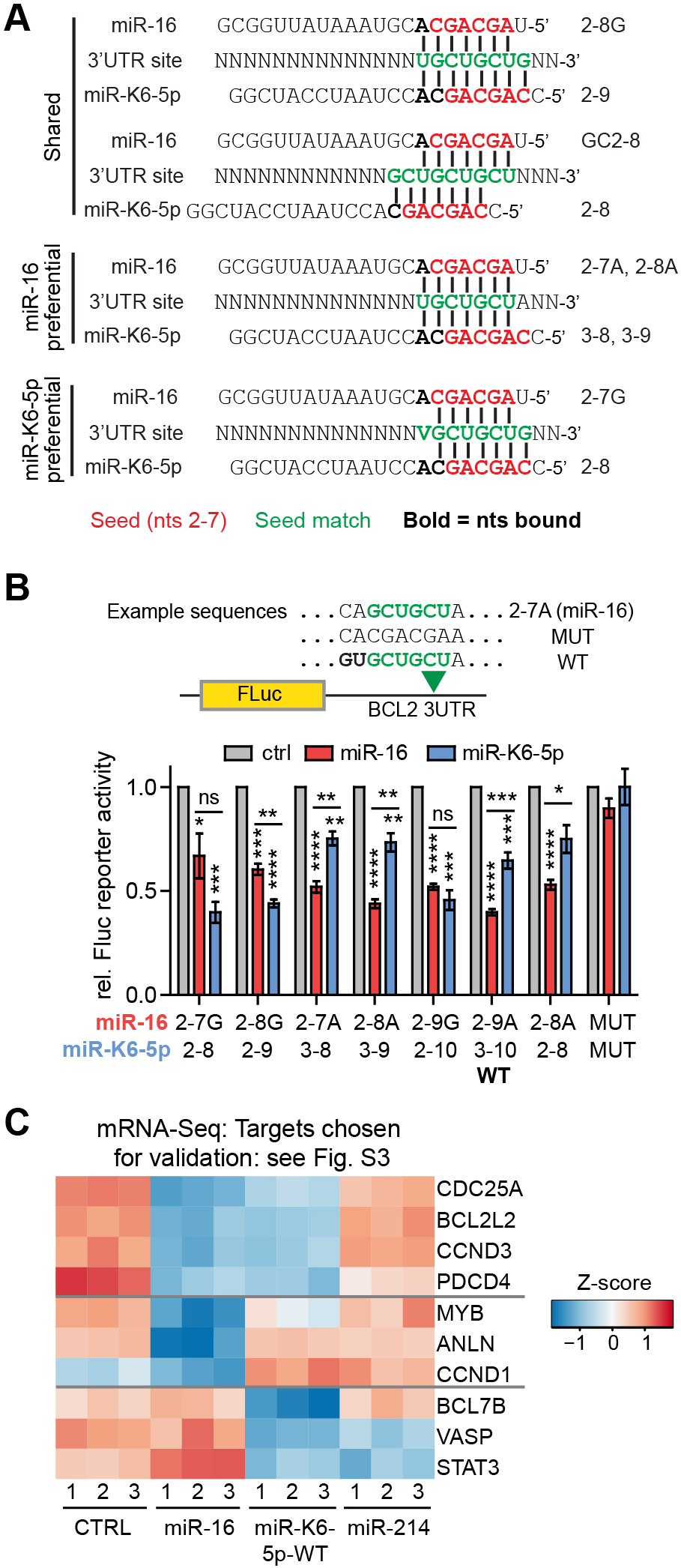
miR-K6-5p regulates target mRNAs through miR-16 binding sites. (A) Diagram of target sites that are shared or preferential based on canonical seed matching for miR-16 and miR-K6-5p. nt 2-7 seed sequences are in red, nts bound by either miRNA in each scenario are in bold, and green denotes the target site. V denotes A,C, or G. (B) The well validated miR-16 binding site in a firefly (FLuc) luciferase 3’UTR reporter vector for BCL2 was modified to various target sites shown in panel A and tested for regulation by miRNA mimics of miR-16 or miR-K6-5p in dual luciferase reporter assays in 293T/NoDice cells. A co-transfected *Renilla* luciferase construct served as an internal control. Fluc data were sequentially normalized to those obtained for the *Renilla* internal control, empty FLuc vector, and negative control mimic (ctrl). The tested sequences can be found in Fig. S3A. (C) Heatmap showing Z-scores for mRNAs that represent different types of targets of miR-16 and/or miR-K6-5p and were chosen for validation by wt and target site mutant 3’UTR reporter assays and Western analyses (see Figure S3B-M).

To directly test regulation of specific types of sites by both miR-16 and miR-K6-5p, we systematically mutated the known miR-16 binding site in the 3’UTR of BCL2 to other types of sites in the context of a luciferase reporter (Fig. 3B, S3A). Results confirmed that the miR-16 2-8G and 2-9G sites were strongly regulated by both miRNAs. In contrast, 2-8A and 2-9A sites for miR-16 were preferentially regulated by miR-16, while the 2-8 seed match for miR-K6-5p (2-7G for miR-16) was strongly regulated only by miR-K6-5p.

To further validate this concept for actual mRNA targets, we chose the top regulated mRNAs that best illustrate these regulatory relationships for validation by 3’UTR dual luciferase reporter assays (Fig. 3C). We specifically included candidates for targets that underlie the tumor suppressive roles of miR-15/16. These validation experiments used wild-type and target site mutant 3’UTRs for known miR-16 targets that are expected to be shared with miR-K6-5p (CDC25A, BCL2L2 (Chang et al., 2007), CCND3, and PDCD4 (Fu et al., 2018), Fig. S3B-E); three known miR-16 targets with preferential regulation by miR-16 over miR-K6-5p (MYB (Chung et al., 2008), ANLN (Lian et al., 2018), and CCND1 (Chen et al., 2008), Fig. S3 F-H); and three candidates for miR-K6-5p targets that are unlikely to be shared by miR-16 (BCL7B, VASP, and STAT3, Fig. S3I-K). STAT3 is a known target of miR-K6-5p (Ramalingam and Ziegelbauer, 2017). Results overall confirmed the predicted regulatory consequences. The observed regulatory relationships were furthermore confirmed for a subset of the encoded proteins using quantitative Western blotting (Fig. S3L-M). Together, these data illustrate that miR-K6-5p substantially mimics miR-16-induced changes in gene expression by regulating an overlapping set of mRNA binding sites. This mimicry specifically extends to several mRNAs encoding proteins with cell cycle regulatory or anti-apoptotic functions, which may underlie the reported tumor suppressive functions of miR-16. Our data finally show that a smaller subset of the observed regulatory consequences and targets are preferential or specific for either miR-16 or miR-K6-5p. Our results so far demonstrate that K6-5p shares a large portion of its binding sites with miR-16, and thereby strongly, but incompletely, mimics miR-16.

### miR-K6-5p confers a competitive disadvantage in the KSHV-transformed PEL cell line BC-3

Finally, we investigated the function of miR-K6-5p in the context of KSHV infection. There are currently no *de novo* infection models that recapitulate KSHV-induced oncogenic transformation, proliferation and survival of human cells. However, KSHV-transformed patient-derived PEL cell lines are a robust model for B cell transformation by KSHV, since they require latent infection by KSHV for their continued proliferation in culture (Godfrey et al., 2005; Guasparri et al., 2004; Wies et al., 2008). We sought to inhibit miR-K6-5p in the PEL cell line BC-3, where miR-K6-5p is expressed at high levels compared to miR-15/16 seed family members (Gottwein et al., 2011, Fig. S4A, B). To achieve lasting inhibition of miR-K6-5p, we designed a miR-K6-5p-specific lentiviral sponge inhibitor (8SK6-5p, Fig. 4A), an approach where repeats of bulged miRNA binding sites trigger miRNA degradation (Ebert et al., 2007). The specificity of 8SK6-5p for miR-K6-5p over miR-16 was established using miRNA sensors in 293T/NoDice cells (Fig. S4C-D). We next transduced either the empty lentivirus or 8SK6-5p into BC-3 cells and selected the top 25% of GFP expressing cells by fluorescence activated cell sorting (FACS), an approach we have previously used to achieve robust inhibition of KSHV miRNAs in PEL cell lines (Gottwein and Cullen, 2010). Three independent cell pools were generated and specific inhibition of miR-K6-5p over miR-16 was confirmed by qRT-PCR and lentiviral sensors for miRNA activity (Fig. 4B-C). As a sensitive readout for altered proliferation upon inactivation of miR-K6-5p, we performed a competitive fitness experiment (Fig. 4D). For this, we mixed matched naïve BC-3 cells with empty vector control or sponge-transduced BC-3 cells at an approximate ratio of 9:1 (naïve:GFP). In this setting, miR-K6-5p-inhibited cells, but not empty vector control cells, significantly outcompeted naïve cells over time (Fig. 4E), suggesting that inhibition of miR-K6-5p in BC-3 results in a competitive advantage. Overall, this result suggests that miR-K6-5p indeed negatively regulates proliferation or survival in the context of KSHV-infected tumor cells.

**Figure 4.**
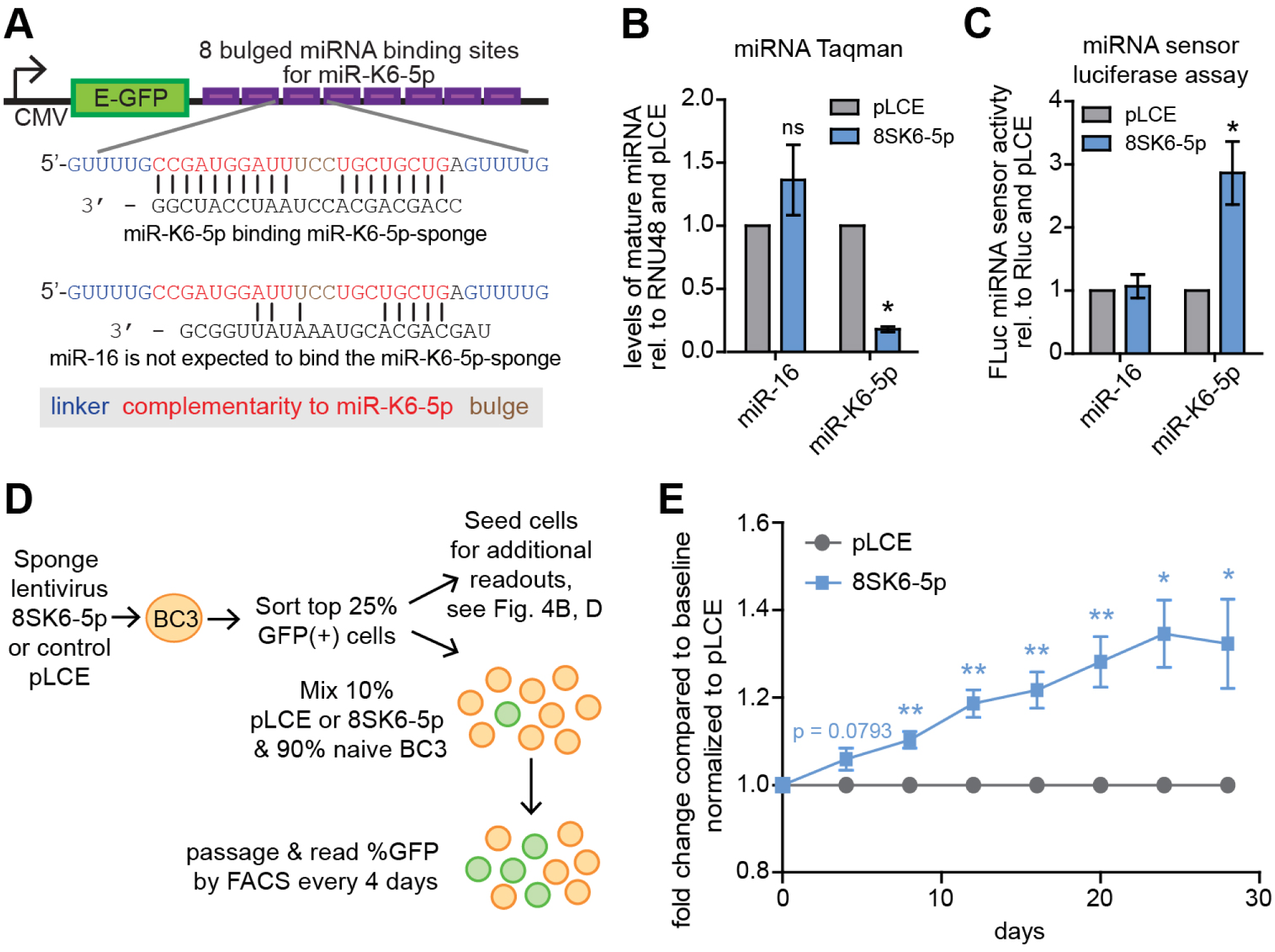
miR-K6-5p confers a competitive disadvantage in the KSHV-transformed PEL cell line BC-3. (A) Diagram of the lentiviral miR-K6-5p-sponge (8SK6-5p). (B) Taqman qRT-PCR was used to assess miR-16 or miR-K6-5p expression in BC-3 cells that were transduced with empty vector (pLCE) or 8SK6-5p (n=3). (C) pLCE or 8SK6-5p-transduced and sorted cells were transduced with luciferase sensor constructs containing three sites of perfect complementarity to miR-K6-5p or miR-16 (n=5). (D) Diagram of experimental design for competition assay in panel (E). (E) Results from competition experiments in BC-3 as outlined in panel (D) and in the text. Data are derived from 3 independent transductions and sorts, 5 independent experiments, and 2 technical replicates per cell line and experiment (n=5 biological replicates). Throughout the figure: * p<0.05, ** p<0.001.

## Discussion

KSHV encodes several viral oncogenes that promote cellular proliferation and survival. These viral oncogenes most likely enable life-long persistence of KSHV infection in the healthy host, but promote tumor formation in the context of immunodeficiency. Here we report that KSHV miR-K6-5p is a mimic of the cellular tumor suppressive miR-15/16 family of miRNAs. Ectopic expression of miR-K6-5p phenocopied miR-16-induced cell cycle arrest. We show that miR-K6-5p phenocopies miR-16-induced changes in cellular mRNA expression and regulates miR-16 target mRNAs through conserved miR-16 binding sites. The mimicry of miR-15/16 by miR-K6-5p extends to targets that are candidates for the mediators of the tumor suppressive roles of miR-15/16, such as the cell cycle regulators CCND3 and CDC25A, and the antiapoptotic protein BCL2L2. Finally, in the KSHV-transformed PEL cell line BC3, inhibition of miR-K6-5p resulted in a significant competitive advantage compared to naïve cells.

Together, our data strongly suggest that miR-K6-5p is an oncoviral mimic of a *bona fide* tumor suppressor. At first glance, it is highly counterintuitive that an oncogenic virus encodes a miRNA with cell cycle inhibitory and tumor suppressive properties. However, since herpesviruses co-evolve with their host, they are exquisitely adapted to establish life-long persistent infection, typically without severe pathogenic consequences in immunocompetent individuals. Thus, viral oncogenesis can be viewed as a non-physiological result of infection that is unlikely to benefit the virus in terms of persistence or transmission (Moore and Chang, 2017). miR-K6-5p is encoded in the major latency locus, where its expression is transcriptionally coupled to that of the major KSHV oncogenes, i.e. LANA, vCyc, vFLIP, and miR-K11. We propose that miR-K6-5p may function as an inbuilt negative regulator to balance the pro-proliferative and pro-survival functions of these oncogenes. Unchecked expression of the viral oncogenes could trigger host responses such as the induction of apoptosis and/or oncogene-induced senescence (Leidal et al., 2012), which would limit viral maintenance. Unchecked viral oncogene expression could also trigger severe pathogenesis, which would eliminate the host. Since miR-15/16 miRNAs are ubiquitously expressed, it is likely that miR-K6-5p has to be highly expressed to boost miR-16-like functionality in the context of infection. Our work suggests that in at least some PEL cell lines (such as in BC-3), miR-K6-5p is expressed at sufficiently high levels to substantially overexpress overall miR-15/16 family miRNA expression. Interestingly, BC-3 cells also express extraordinarily high levels of vCYC (Jarviluoma et al., 2004, see also Fig. S4B). A recent report suggests that the intronic region encoding most of the KSHV miRNAs, including miR-K6-5p, is also highly expressed in KS lesions (Rose et al., 2018).

In addition to its overexpression of miR-15/16-like functions, miR-K6-5p may have other advantages for the virus. The expression of miR-15/16 family miRNAs is cell cycle regulated (Rissland et al., 2011). It is possible that miR-K6-5p escapes similar cell cycle regulation and thereby differs from miR-15/16 family miRNAs, although this remains to be tested. Finally, miR-K6-5p has evolved at least some targets that differentiate its activity from that of the miR-15/16 family miRNAs and introduce unique regulatory interactions into infected cells (for example the regulation of STAT3, see Ramalingam and Ziegelbauer, 2017). This is in keeping with the altered functionality of tumor viral oncogenes over their cellular counterparts. While this study presents the first evidence of a virus specifically mimicking a known cellular tumor suppressor, the tumor virus Epstein-Barr virus encodes a viral protein (EBNA3B) that reduces the oncogenic potential of the virus in mice (White et al., 2012). However, this tumor suppressive function of EBNA3B is thought to be due to increased recruitment of T cells to EBV-infected B cells, while miR-K6-5p acts in a direct, cell intrinsic manner.

All together, we show that the oncogenic herpesvirus KSHV encodes a mimic of the cellular tumor suppressive miR-15/16 miRNA family. This finding highlights the complexity of KSHV-mediated regulation of host gene expression, which must balance the need to maintain a latent viral reservoir with avoiding a detrimental pathogenic outcome that would limit viral persistence.

## Acknowledgements

We would like to thank Drs. Lauren Gay and Rolf Renne (University of Florida) for qCLASH data files, Hal Bogerd and Dr. Bryan Cullen (Duke University Medical Center) for 293T/NoDice, the University of Chicago Genomics Facility for Illumina sequencing, the Immunobiology Center Flow Cytometry Core and the Robert H. Lurie Comprehensive Cancer Center Flow Cytometry Core Facility for FACS, and Neil Kuehnle for comments on the manuscript. This study was supported by the National Cancer Institute grant R01 CA180813, by ACS Research Scholar Grant RMC-129546, by Searle and Zell Scholar Awards from the Robert H. Lurie Comprehensive Cancer Center to E.G. K.M. was funded by the Cellular and Molecular Basis of Disease Training Program (T32 GM008061). The content is solely the responsibility of the authors and does not necessarily represent the official views of the funding agencies.

## Author Contributions

K.M., M.M., and E.G. designed the study, analyzed, and interpreted the data; K.M. and E.G. wrote the manuscript; K.M. conducted all experiments, except for Fig. S1A (E.G) and S4B (M.M); M.M derived 293T/DKO; K.M. constructed most vectors, several vectors were made by M.M. or K.C; M.S. provided initial analysis of mRNA-Seq data and 3’UTR file; E.T.B. contributed the pipeline used for miRNA seed matching.

## Methods

### Cell culture

Primary lymphatic endothelial cells (LEC, PromoCell, adult donor 419Z035.4) were maintained in Endothelial Cell Basal Medium MV2 supplemented with EGM-2 bullet-kit (PromoCell) and used before passage 7. 293T/NoDice and 293T/DKO cells were maintained in Dulbecco modified Eagle medium (DMEM) containing 4.5 g/liter glucose and L-glutamine (Corning), supplemented with 10% Serum Plus-II (Sigma-Aldrich). The PEL cell line BC3 was maintained in RPMI 1640 medium containing L-glutamine (Corning), supplemented with 0.05mM β-mercaptoethanol (Sigma-Aldrich) and 20% Serum Plus-II (Sigma-Aldrich).

### Cloning procedures

Oligonucleotides and dsDNA fragments used in cloning procedures are listed in Table S4. Lentiviral miRNA expression vectors were based on the published vector pLCE (Zhang et al., 2009). To clone the miR-16-1 expression vector, a 250bp fragment centered on the miR-16-1 pre-miRNA stem-loop was amplified from genomic DNA from the PEL cell line BC-1, using the primers 1559 and 1560, which add XhoI and NotI cloning sites. Because the natural pre-miR-K6 stem-loop expresses higher levels of miR-K6-3p than miR-K6-5p, we designed a strategy to express only miR-K6-5p from the miR-122 stem-loop, which is naturally processed to produce high levels of the mature 5p miRNA and with almost no expression of the 3p arm. We re-engineered pri-miR-122 to replace the miR-122-5p sequence miR-K6-5p, while maintaining the predicted secondary structure of the stem-loop (Fig. S2B). A dsDNA fragment encompassing a 250 bp hybrid pri-miR-122/K6-5p sequence and flanking XhoI and NotI cloning sites was ordered from Epoch Life Science (Missouri City, TX), cloned in the intermediate vector pBSK. To simultaneously delete the human miR-15a/16-1 and miR-15b/16-2 loci in 293T/DKO, we employed the double nicking strategy by Cas9 D10A to avoid off-target effects (68). Specifically, we designed 8 sgRNAs, with 2 sgRNAs on each side of the miR-15a/16-1 and miR-15b/16-2 loci. The relevant oligonucleotides were annealed and cloned into BbsI-digested (Fermentas) pX335, which encodes the Cas9 D10A nickase mutant. Full-length wild-type 3’UTR sequences were amplified from PEL cell line genomic DNA, and cloned into the published firefly luciferase reporter construct pL/SV40/Fluc (pLSG, Gottwein et al., 2007), using XhoI and NotI restriction sites. For 3’UTR reporter mutants, miRNA target sites were mutated individually and in combination using PCR with mutant primers and inserted between the XhoI and NotI restriction sites of pLCE using Gibson assembly (Gibson, 2011). 8SK6-5p was based on the previously described lentiviral vector pLCE (Zhang et al., 2009) and constructed by annealing of 2 pairs of ultramer oligonucleotides (Integrated DNA Technologies) and insertion between XhoI and XbaI sites in the 3’UTR of pLCE by three-fragment ligation. For cleavage sensors of miRNA activity (pLCG-3T16 and pLCG-3TK6, Figs. 4B, S4B-D), oligonucleotides were designed and annealed to create a sequence containing 3 sites of perfect complementarity to miR-K6-5p and inserted between Xho1 and Xba1 sites of the published lentiviral reporter vector pL/CMV/Fluc (pLCG, Linnstaedt et al., 2010). Complete sequences of all inserts were verified by Sanger sequencing.

### Generation of HEK293T miR-15/16 double knockout (293T/DKO) cell line

The eight sgRNA constructs described above were co-transfected in 293T cells. After 2 days, cells were single-cell cloned into 96-well plates at a calculated 0.25 cells/well. Cell clones were PCR-screened for homozygous deletions using the primer pairs 1824/1825 (for miR-15a/miR-16-1) and 1903/1909 (for miR-15b/miR-16-2). Loss of mature miR-16 expression was confirmed from clones positive for homozygous deletions by primer extension and Taqman miRNA assay.

### Luciferase reporter assays

For miRNA binding site validation in 293T/NoDice cells, cells were seeded at 200,000 cells per well in 24-well plates. The next day, cells were co-transfected with 2.5ng/ml pLSG (empty vector, or modified to include wt or mutant test 3’UTRs); 5ng/ml of the internal control pL/SV40/Rluc (pLSR) *Renilla* luciferase construct (Gottwein et al., 2007); 5pmol of negative control #1, miR-16, or miR-K6-5p mimics (miRVana); and 0.3 μg of pLCE as a DNA carrier. Cells were transfected using lipofectamine 2000 as instructed by manufacturer (Life Technologies). Two days later, growth medium was aspirated and cells were lysed in 1X passive lysis buffer (Promega) without washing. 5 uL of lysate from each condition were loaded into a 96-well half-area well plate (Greiner) and dual luciferase assays were performed using Promega Dual Luciferase Kit on a Victor Nivo plate reader. Firefly activity was normalized to *Renilla* luciferase from the same well and resulting ratios were further normalized to those from wells that had received the empty pLSG vector and the control mimic. Reporter assays shown in Fig. S4C-D were done similarly, except that 0.4 nM of either control, miR-16, or miR-K6-5p mimic (miRVana) were used. For lentiviral reporter assays in sorted BC-3 cells, cells were seeded at 250,000 cells/ml in 24-well plates and co-transduced with lentiviruses pLCG (empty vector control), pLCG-3T16, or pLCG-3TK6 and pL/CMV/RLuc (pLCR, *Renilla* luciferase internal control, Linnstaedt et al., 2010). Two days later, cells were pelleted, lysed in 100 uL 1X passive Lysis Buffer (Promega) and processed for dual luciferase reporter assays as above. Firefly luciferase activities were normalized for *Renilla l*uciferase activities. The data were further normalized to those obtained using the empty pLCG vector and pLCE-transduced cells. Statistical analyses were performed using paired, two-tailed Student’s t-tests.

### Western blot analysis

293T/NoDice cells were transfected with 10 nM concentrations of control, miR-16, or miR-K6-5p mimic using lipofectamine RNAiMax as instructed by manufacturer. 2 days later, cells were harvested for Western blot analysis. Culture medium was aspirated, cells were washed with 1X Phosphate Buffered Saline (PBS) (Corning), and scraped into 100 uL ice-cold RIPA lysis buffer containing 1X protease inhibitor cocktail III (Calbiochem, EMD Millipore, Darmstadt, Germany) and 1X PhosSTOP phosphatase inhibitor cocktail (Roche, Mannheim, Germany). Cells were briefly vortexed and then lysed on ice for 15 minutes. Lysates were sonicated for 30 second intervals (30 seconds on, 30 seconds off) for 7 cycles of sonication in a 4°C water bath using the Bioruptor Sonication System (Diagenode, Denville, NJ) at the high-intensity setting. Sonicated lysates were cleared by centrifugation at 16,000 x g for 15 minutes at 4°C. Lysates were subsequently quantified using ThermoFisher BCA kit. NuPAGE LDS sample buffer was added to a final concentration of 1X (ThermoFisher Scientific), samples were heated at 70°C for 10 minutes. Equal protein amounts were run on 4-12% Bis/Tris gels in 1X NuPAGE MOPS SDS Running Buffer (ThermoFisher Scientific). Proteins were transferred to nitrocellulose membranes, which were blocked using 5% non-fat milk powder in 1X PBS for 1 hour at room temperature. Primary antibodies are listed in Table S4 and were used at indicated dilutions, in 5% non-fat milk powder in 1X PBS and 0.2% Tween20 and incubated overnight at 4°C. Membranes were washed 3 times in 1X PBS containing 0.2% Tween20. Primary antibodies were detected with IRDye 800 CW-conjugated goat anti-rabbit or anti-mouse IgG secondary antibodies (LI-COR Biosciences, Lincoln, NE), diluted 1:20,000 in 1X PBS, 1 hour RT), and imaged with the Odyssey Fc Dual-Mode Imaging System (LI-COR). Results were quantified using ImageStudio and compared statistically using using paired, two-tailed Student’s t-tests.

### Growth curve and cell cycle analyses

Cells were transfected with 10 nM control, miR-16, or miR-K6-5p mimic (miRVana) with Lipofectamine RNAiMax (ThermoFisher Scientific, Cat No: 13778150) as instructed. Cells were counted manually and split to equal numbers every other day for 8-14 days, as indicated. Total cell numbers were calculated using dilution factors from the previous days, and normalized to the total cell number for control-transfected cells. Propidium iodide (PI) staining of LEC was performed two days after transfection. LEC were trypsinized, washed once with 1X PBS and fixed at −20°C overnight using 70% ethanol. Fixed cells were pelleted, washed three times with 1X PBS, resuspended and stained for 30 minutes in propidium iodide (PI)/RNase staining buffer (BD Pharmingen). PI-stained cells were subjected to flow cytometry and resulting data were analyzed by FlowJo, by fitting to the Watson (Pragmatic) cell cycle model. Cell cycle analysis in 293T/NoDice by PI/anti-BrdU staining was performed two days after transfection. For this, transfected cells were pulsed with 75μM bromodeoxyuridine (BrdU) for 45 minutes. Cells were then trypsinized, washed with 1X PBS, and fixed at −20°C overnight using 70% ethanol. Fixed cells were pelleted, washed with 1X PBS three times, and then incubated with 1.5M HCl for 30 minutes to denature DNA. Cells were washed 3 times in PBS, incubated with FITC-anti-BrdU (BD Biosciences) for 30 minutes at room temperature in the dark, washed once with PBS, resuspended in PI/RNase staining buffer (BD Pharmingen) and incubated in the dark at room temperature for 30 minutes. PI and anti-BrdU co-stained cells were analyzed by flow cytometry on a FACS Canto II and resulting data were analyzed in FlowJo.

### Lentivirus production for sponge-transduction experiments

pLCE-based vectors were produced by co-transfection with pMDLgpRRE, pRSV-Rev, and pVSV-G into 293T cells via polyethylenimine HCl MAX (PEI, Polysciences, Cat No: 24765) diluted in Opti-MEM (Gibco, Cat No: 11058021) at a ratio of PEI:DNA = 3uL (stock solution 15.6mM) :1ug. Medium was exchanged to BC-3 growth medium 6 hours after transfection. 72 hours after transfection, supernatants were harvested, centrifuged to remove debris, filtered through 0.45μm syringe filters, and concentrated using 100kDa cutoff Amicon Ultra-15 centrifugal filters (Fisher, Cat No: UFC910024). Resulting concentrated viruses were titrated in BC3 cells by serial dilutions and measuring %GFP(+) cells by FACS.

### Lentiviral sponge transduction, sort, and competition assay

BC-3 cells were seeded at 250,000 cells/ml and transduced with pLCE or pLCE-8SK6-5p at a multiplicity of infection (MOI) of 5, in the presence of 4 μg/ml polybrene. Two days after transduction, cells were sorted on a FACS Aria to collect the top ∼25% GFP-expressing cells (RHLCC Flow Cytometry Core, Northwestern University). Collected cells were seeded at 500,000 cells/ml and allowed to recover and expand for several days before functional assays. We derived three independent cell pools by individual transductions and sorts. For competition assays, sorted GFP(+) cells from each transduction were counted and mixed with matched naïve BC3 cells at a ratio of approximately 1:10 (10% GFP(+) and 90% naïve BC-3). Cell mixes were subjected to flow cytometry to determine the baseline %GFP for day 0 and subsequently split to 250,000 cells/ml every other day, and the percentage of GFP(+) cells was measured by flow cytometry every 4 days for 28 days. Two technical replicates per mixed populations were completed for each of five biological replicates (n=5, including 2 replicates from sort 1 and 3 and one replicate from sort 2). Data from technical replicates were averaged before analysis of biological replicates. For the analysis of biological replicates, data were sequentially normalized to baseline % of GFP+ cells and then to data from the pLCE control.

### RNA preparation and miRNA quantification by qRT-PCR

Cells were resuspended in TRIzol (ThermoFisher Scientific, Cat No: 15596018) and RNA was isolated using the Direct-zol RNA MiniPrep Plus kit (Zymo Research, Cat No R2070), omitting the DNase treatment step. For mRNA-Seq, total RNA was submitted to the University of Chicago Genomics core facility. For quantitative real-time PCR (qRT-PCR), total RNA was treated with RQ1 DNase for 30 minutes at 37°C (Promega, Cat No: PAM6101). The DNase reaction was stopped with RQ1 Stop Solution for 10 minutes at 65°C (Promega). qRT-PCR was performed on 5 ng of total RNA using TaqMan mature miRNA assays and an RNU48 control (Life Technologies). Real time PCR was performed on a Roche LightCycler® 480 system and data were analyzed using the double delta CT method, with results from hsa-miR-16 and kshv-miR-K12-6-5p expression normalized to those from the control RNU48.

### Bioinformatics

For mRNA-Seq analysis, the quality of DNA reads, in fastq format, was evaluated using FastQC. Adapters were trimmed and the reads were aligned to the human genome (hg19) using STAR (Dobin et al., 2013). Read counts for each gene were calculated using htseq-count (Anders et al., 2015) in conjunction with a gene annotation file for hg19 obtained from Ensembl GRCh37.75 (http://useast.ensembl.org/index.html). Normalization and differential expression were determined using DESeq2 (Love et al., 2014). For all bioinformatics analysis, only genes that had measurable expression in all samples were considered. Pearson correlation coefficients were calculated from pairwise comparisons of log_2_ fold changes using *pandas.* GSEA was conducted on gene lists that were ranked by fold change using GSEA preranked and the classic enrichment statistic under default settings. For 3’UTR seed matching, annotated 3’UTRs were extracted from the human genome using the BioMart tool on Ensembl GRCh37.75 (http://useast.ensembl.org/biomart/martview/db174d84ffc0353f3fa25c30697cd99a). A custom perl script was used to collect all annotated 3’ UTRs for each gene. For genes with more than one annotated 3’ UTR, the longest 3’UTR was kept for analysis. miRNA binding sites were identified using a custom pipeline available on Code Ocean (https://doi.org/10.24433/CO.1360162.v1). The analyzed mRNA-Seq dataset can be found in Table S1 and raw data are available in GEO (GSE128576). Targetscan predictions for miR-15/16, miR-214, miR-34, miR-1, miR-155, and miR-21 were downloaded from the Targetscan 7.1 website, sorted from low to high cumulative weighted context++ scores and filtered for mRNAs that are designated as expressed in the mRNA-Seq dataset and have annotated Ensembl 3’UTRs according to our analysis above. Only 3’UTR targets were considered. The top 250 remaining predicted targets for each miRNA were selected and used for CDF plot analyses and 2-sample Kolmogorov–Smirnov (K-S) tests, which were done using Biopython. DAVID pathway analysis was performed on resulting top 250 predicted miR-15/16 targets using all human genes as background.

### Data and Software Availability

Custom Code is available on Code Ocean (https://doi.org/10.24433/CO.1360162.v1). The analyzed mRNA-Seq dataset can be found in Table S1 and raw data are available in GEO (GSE128576).

## Supplementary Tables

### Table S1 Analyzed 293T/NoDice mRNA-Seq dataset. Related to Figs. 2, S2

(A) Differential gene expression control vs. miR-16 mimics.

(B) Differential gene expression control vs. miR-K6-5p mimics.

(C) Differential gene expression control vs. miR-K6-5’U mimics.

(D) Differential gene expression control vs. miR-214 mimics.

(E) Summary of differential gene expression results, including miRNA target information from simple matching, PAR-CLIP, qCLASH, and predictions by Targetscan 7. Columns indicate Gene Symbol (Gene), LOG2 fold changes (LOG2FC), linear fold changes (LinFC), and false discovery rate-adjusted p values (FDRadjP) for miR-16 (M16), miR-K6-5p (K6WT), miR-K6-5p-5’U (K6U), or miR-214 (M214). The column CDF_incl specifies the 16,546 genes that were included in CDF analyses in Fig. 2D-F and Fig. S2. “PAR-CLIP_miR-16” specifies 3’UTR seed targets for miR-16 as detected in the published PAR-CLIP dataset (see Fig. S2E, Gottwein et al., 2011), qCLASH_miR-16 specifies 3’UTR seed targets for miR-16 as detected in the published qCLASH dataset (see Fig. S2E, Gay et al., 2018). Columns TS1516ALL, TS1516TOP250, TS214_TOP250 specify genes that are Targetscan7.1-predicted 3’UTR targets of miR-15/16 family miRNAs, or belong to the top 250 predicted targets of miR-15/16 family miRNAs or miR-214, respectively, and were analyzed in Figs. 2D-F or Fig. S2E). Columns M16_2to8 and M16_2to8A specify genes included in Fig. S2E. Columns V to AS specify genes that contain indicated seed matches and are analyzed in Fig. S2F-I. Columns AK to AW include complete predictions for Targetscan 7.2 for miR-15/16.

**Table S2.** DAVID Pathway analysis of the Top250 Targetscan predicted mRNA targets of miR-15/16 family miRNAs. Genes that contribute to the observed enrichments in cell-cycle related terms were selected and included in the heatmap shown in Fig. 2G.

(A) Analysis for biological pathways, i.e. “GOTERM_BP_DIRECT”.

(B) Analysis for molecular functions “GOTERM_MF_DIRECT”.

(C) Analysis for KEGG categories.

**Table S3.** qCLASH clusters for detected for miR-K6-5p in any of the three published replicates (Gay et al., 2018).

**Table S4.** Oligonucleotides (A) and antibodies (B) used in this study.

