## Supplementary Figures for "The oncogenic Kaposi’s sarcoma-associated herpesvirus encodes a mimic of the tumor suppressive miR-15/16 miRNA family"

### Figure S1

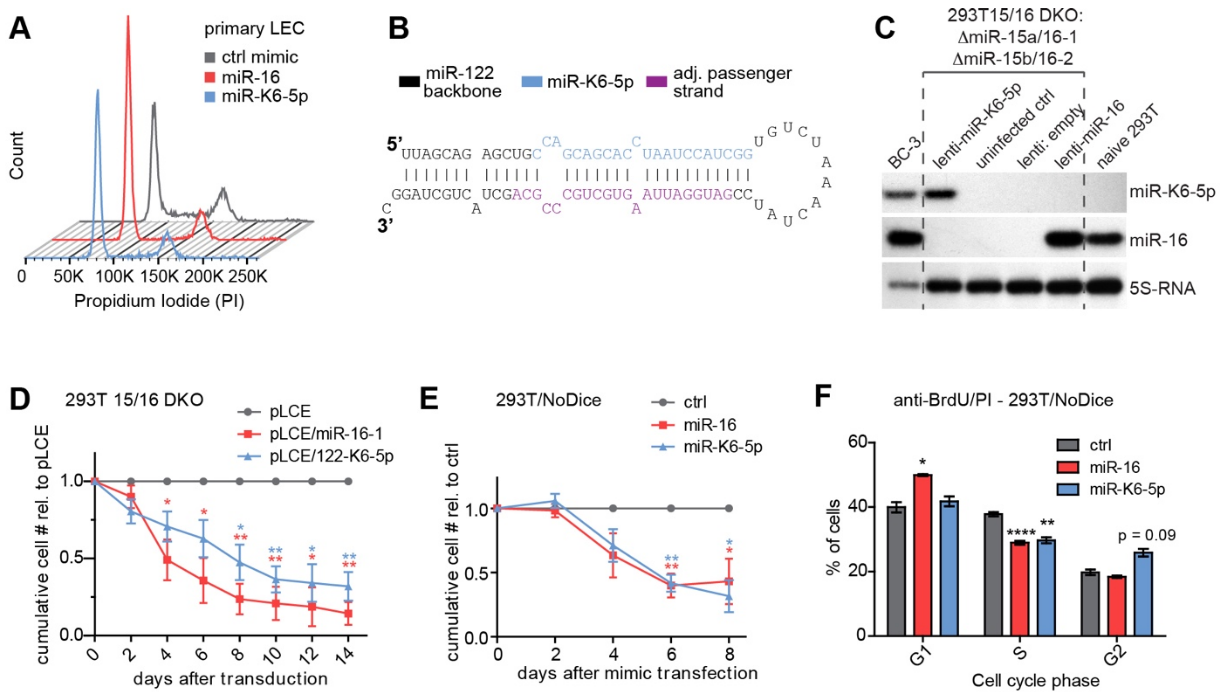


**Figure S1. Related to Fig. 1.**

(A) Representative example of the data used to generate Fig. 1C.

(B) Schematic of the stem-loop structure used for lentiviral expression of miR-K6-5p from the miR-122 precursor (see panels C, D).

(C) 293T with CRISPR-mediated deletion of both miR-15/16 loci (293T15/16 DKO) were transduced with lentiviral constructs expressing miR-16, miR-K6-5p, or empty vector. Primer extension analysis verified complete loss of miR-16 expression in 293T15/16 DKO as well as correct 5'end processing and relatively physiological expression of the lentivirally expressed miRNAs.

(D) 293T15/16 DKO cells stably transduced with the vectors described in (C) were seeded at equal numbers and split to equal numbers every other day for 14 days. Cell numbers were normalized to those from pLCE-transduced cells (n=3)

(E) 293T/NoDice cells were seeded at equal numbers and transfected with miRNA mimics of miR-16, miR-K6-5p, or a negative control (ctrl) mimic. Cells were counted every 2 days and split to equal cell numbers over the course of 8 days. Cell numbers were normalized to those of ctrl-transfected cells. (n=4).

(F) 293T/NoDice were transfected as described for (E). 4 days after transfection, cells were pulsed with 75uM BrdU for 45 min, then harvested and fixed. Fixed cells were then stained with propidium iodide (PI) and a FITC-labeled anti-BrdU antibody to assess DNA content and percentage of cells in S phase, respectively. FITC/PI co-stained cells were analyzed by flow cytometry, and the resulting data were analyzed for cell cycle stage using FlowJo. (n=3). Throughout: * $p<0.05$, ** $p<0.01$, *** $p<0.001$, **** $p<0.0001$, error bars s.e.m.

**Figure S2**

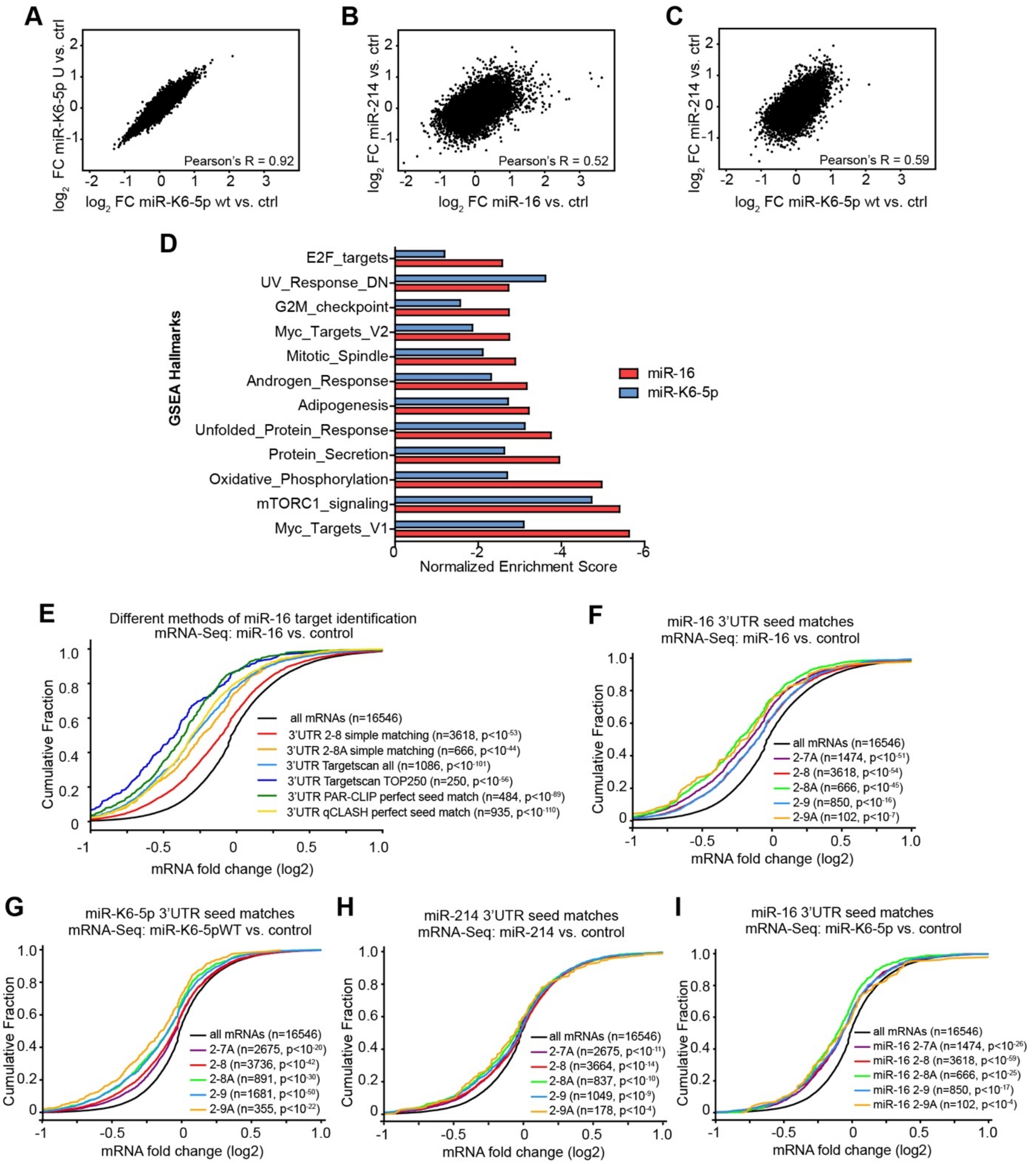


**Figure S2. Related to Fig. 2.**

(A-C) Comparison of log2 mRNA fold changes in cells transfected with miR-K6-5p wt or miR-K6-5p-U (A), in cells transfected with miR-16 or miR-214 (B), and in cells transfected with miR-K6-5p wt or miR-214 (C).

(D) Gene set enrichment analysis (GSEA) identifies enrichments of miR-16- or miR-K6-5p-downregulated genes in categories that are broadly associated with cellular survival and proliferation, among others. All hallmark categories that were enriched in either miR-16 or miR-K6-5p-downregulated mRNAs with a normalized enrichment score of -2.50 or less were included in the graph.

(E) Cumulative distribution frequency (CDF) analyses of the miR-16 mRNA-Seq data were performed to establish a robust set of miR-16 targets for analysis. We compared candidates for miR-16 targets that were predicted by Targetscan v7.1 (Agarwal et al., 2015), including all 3'UTR targets or those that ranked in the top 250, simple matching for the presence of 2-8 or 2-8A 3'UTR seed matches of miR-16, a published PAR-CLIP dataset (Gottwein et al., 2011), and a published qCLASH dataset (Gay et al., 2018). *p* values of comparisons to all mRNAs were calculated using 2-sample K-S tests.

(F-H) CDF analyses were performed to establish regulation of mRNAs with canonical 3'UTR seed matches for miR-16 (F), miR-K6-5p (G), or miR-214 (H) in mRNA-Seq by the individual targeting miRNAs. *p* values of comparisons to all mRNAs were calculated using 2-sample K-S tests.

(I) CDF analyses were performed to test for regulation of mRNAs with canonical 3'UTR seed matches for miR-16 in miR-K6-5p-transfected 293T/NoDice. *p* values of comparisons to all mRNAs were calculated using 2-sample K-S tests.

See Table S1 for genes included in the gene sets shown in panels E-I.

**Figure S3.**

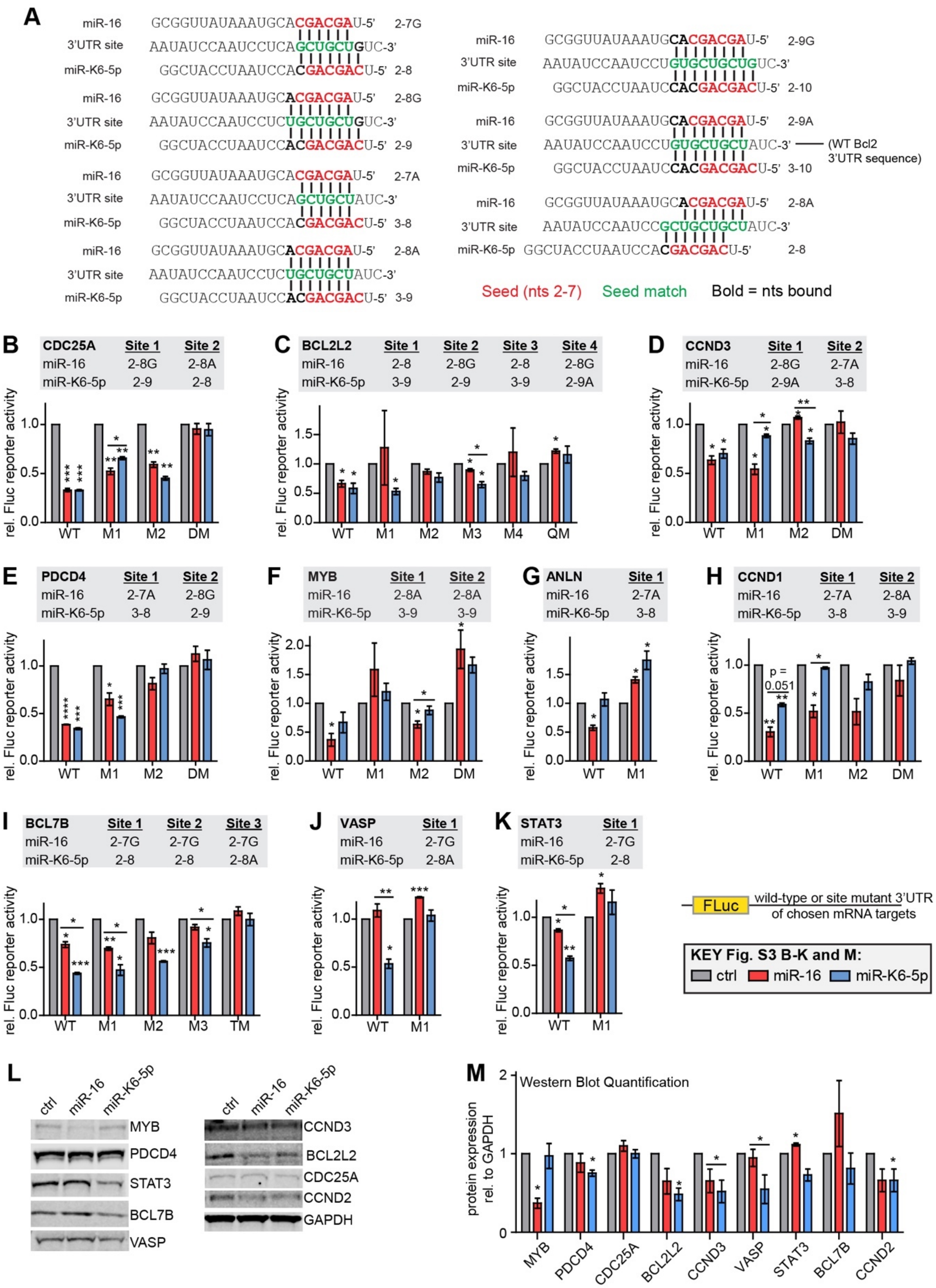

A
miR-16 GCGGUUAUAAAUGCACGACGAU-5' 2-7G
3'UTR site AAUAUCCAAUCCUCAGCUGCUGUC-3'
miR-K6-5p GGCUACCUAAUCCACGACGACU-5' 2-8
Seed (nts 2-7)
Seed match
Bold = nts bound
(WT Bcl2 3'UTR sequence)
B CDC25A Site 1 Site 2
C BCL2L2 Site 1 Site 2 Site 3 Site 4
D CCND3 Site 1 Site 2
E PDCD4 Site 1 Site 2
F MYB Site 1 Site 2
G ANLN Site 1
H CCND1 Site 1 Site 2
I BCL7B Site 1 Site 2 Site 3
J VASP Site 1
K STAT3 Site 1
rel. Fluc reporter activity
FLuc
wild-type or site mutant 3'UTR of chosen mRNA targets
KEY Fig. S3 B-K and M:
ctrl
miR-16
miR-K6-5p
L
MYB
PDCD4
STAT3
BCL7B
VASP
CCND3
BCL2L2
CDC25A
CCND2
GAPDH
M
Western Blot Quantification
protein expression rel. to GAPDH

**Figure S3. Related to Fig 3.**

(A) Sequences of the BCL2-3'UTR seed match mutants that were tested for regulation by miR-16 or miR-K6-5p using dual 3'UTR luciferase reporter assays in Fig. 3B.

(B-K) Wild-type and miR-16 and/or miR-K6-5p binding site mutant 3'UTR luciferase reporter vectors for CDC25A (B), BCL2L2 (C), CCND3 (D), PDCD4 (E), MYB (F), ANLN (G, CCND1 (H), BCL7B (I), VASP (J), and STAT3 (K) were tested for regulation by miR-16 and miR-K6-5p in 293T/NoDice cells. M: mutant, DM: double mutant, TM: triple mutant, QM: quadruple mutant. Data from Fluc reporters were sequentially normalized to those from a co-transfected *Renilla* luciferase control and values obtained by transfection with negative control mimic. n=3.

(L) Representative Western blot analyses of 293T/NoDice transfected with ctrl, miR-16 or miR-K6-5p. Cell lysates were collected 2 days after transfection and analyzed for expression of the indicated proteins using quantitative Western blotting on the Licor platform.

(M) Quantification of data from (M) (n=3).

Throughout the figure: * $p<0.05$, ** $p<0.01$, *** $p<0.001$, **** $p<0.0001$.

**Figure S4**

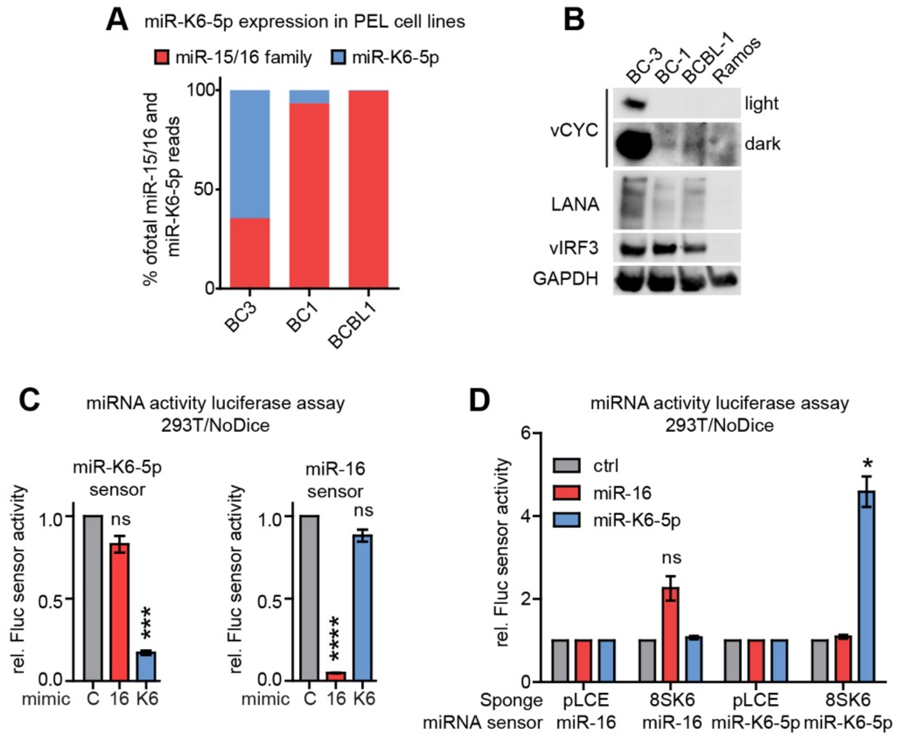


**Figure S4. Related to Fig.4.**

(A) Proportion of small RNA reads for miR-15/16 family members (i.e. the cellular miRNAs included in Fig. 1A, except miR-214) and miR-K6-5p in small RNA sequencing datasets from the PEL cell lines BC-3, BC-1, and BCBL-1 (Gottwein et al., 2011).

(B) Quantitative Western blot for KSHV latent proteins vCYC, LANA, and vIRF3, in the PEL cell lines BC-3, BC-1, BCBL-1, and the KSHV-negative B cell line Ramos. GAPDH is a loading control.

(C) miRNA sensors for miR-K6-5p or miR-16 specifically report the activity of the intended miRNA, with minimal cross-reporting. 293T/NoDice were co-transfected with 0.4 nM of ctrl, miR-16, or miR-K6-5p mimic and Firefly luciferase sensors with three sites of perfect complementarity to miR-K6-5p (left) or miR-16 (right), or an empty Fluc control vector (pLCG). Data obtained for each miRNA sensor were sequentially normalized to those obtained with a cotransfected internal Renilla luciferase control vector (pLCR), the control mimic and pLCG (n=3).

(D) The miR-K6-5p sponge (8SK6-5p) is specific for miR-K6-5p over miR-16. 293T/NoDice were co-transfected with plasmids expressing the empty vector (pLCE) or pLCE/8SK6-5p and 0.4 nM of ctrl, miR-16, or miR-K6-5p mimics as well as the perfect miRNA sensor plasmids for miR-16 (3T16), miR-K6-5p (3TK6), or the control vector (pLCG). All samples also received the internal control plasmid pLCR. Data obtained for each miRNA sensor were sequentially normalized to those obtained with pLCR, the control mimic and pLCG (n=3).
